# Ripening dynamics revisited: an automated method to track the development of asynchronous berries on time-lapse images

**DOI:** 10.1101/2023.07.12.548662

**Authors:** Benoit Daviet, Christian Fournier, Llorenç Cabrera-Bosquet, Thierry Simonneau, Maxence Cafier, Charles Romieu

**Affiliations:** LEPSE, Univ Montpellier, INRAE, Institut Agro, Montpellier, France; AGAP Institut, Univ Montpellier, CIRAD, INRAE, Institut Agro, Montpellier, France

**Keywords:** High-throughput phenotyping, Computer vision, Grapevine berry, Fruit detection, Fruit segmentation, Tracking

## Abstract

**Background:** Grapevine berries undergo asynchronous growth and ripening dynamics within the same bunch. Due to the lack of efficient methods to perform sequential non-destructive measurements on a representative number of individual berries, the genetic and environmental origins of this heterogeneity, as well as its impacts on both vine yield and wine quality, remain nearly unknown. To address these limitations, we propose to track the growth and coloration kinetics of individual berries on time-lapse images of grapevine bunches.

**Result:** First, a deep-learning approach is used to detect berries with at least 50±10% of visible contours, and infer the shape they would have in the absence of occlusions. Second, a tracking algorithm was developed to assign a common label to shapes representing the same berry along the time-series. Training and validation of the methods were performed on challenging image datasets acquired in a robotised high-throughput phenotyping platform. Berries were detected on various genotypes with a F1-score of 91.8%, and segmented with a mean absolute error of 4.1% on their area. Tracking allowed to label and retrieve the temporal identity of more than half of the segmented berries, with an accuracy of 98.1%. This method was used to extract individual growth and colour kinetics of various berries from the same bunch, allowing us to propose the first statistically relevant analysis of berry ripening kinetics, with a time resolution lower than one day.

**Conclusions:** We successfully developed a fully-automated open-source method to detect, segment and track overlapping berries in time-series of grapevine bunch images. This makes it possible to quantify fine aspects of individual berry development, and to characterise the asynchrony within the bunch. The interest of such analysis was illustrated here for one genotype, but the method has the potential to be applied in a high throughput phenotyping context. This opens the way for revisiting the genetic and environmental variations of the ripening dynamics. Such variations could be considered both from the point of view of fruit development and the phenological structure of the population, which would constitute a paradigm shift.

## Background

Unlike climacteric fruits (e.g. bananas, apples or mangoes) which accumulate sufficient starch reserve to achieve post-harvest ripening, the grape berry necessarily ripens on the vine, at the rate of translocation of water and sucrose through the phloem. Ripening involves the sudden activation of the apoplastic pathway of phloem unloading [Zhang et al., 2006], which leads to the second growth phase during which each berry accumulates about 1M hexoses and becomes coloured [Ojeda et al., 1999; Houel et al., 2013; Bigard et al., 2022; Shahood et al., 2020]. Then, the definitive stop of phloem unloading triggers a more or less pronounced shrivelling period known as overripening [Savoi et al., 2021; McCarthy 1999]. It is widely accepted that these dynamic processes are under strong developmental and transcriptomic control [Rienth et al., 2016; Fasoli et al., 2018], and may vary according to the genotype and its interaction with environmental conditions (GxE), particularly light, temperature, and water availability [Suter et al., 2021]. A considerable research effort is devoted to understanding the phenological, physiological and molecular origins of global change effects on grapevine yield and quality [Pastore et al., 2022], which therefore requires dynamic monitoring of the ripening process.

Due to the lack of efficient non-destructive phenotyping methods to study berries individually, the body of knowledge is mainly based on measurements of the average evolution in periodic samples of randomly selected berries. This approach overlooks the heterogeneity of the fruit population representative of the future harvest. Moreover, scarce studies on single berries recently revealed how chimerical these population-relevant samples are, both on the phenology and metabolic point of view. For example, the fact that the developmental lag between two berries can be almost as long as their growth duration leads to a two fold overestimation of the duration of the second growth period, when considering the average evolution among several berries [Shahood et al., 2020; Bigard et al., 2022]. Furthermore, mixing growing and shrivelling berries leads to the averaging bias that a constant volume is maintained during late ripening, and that excess water from the phloem mass flow must be released into the xylem backflow [Keller et al., 2015]. There is thus an urgent need to develop methods for temporal and non-destructive monitoring of cohorts of individual fruits in the future harvest.

Non-destructive spectrometric methods such as NIR, fluorescence and hyperspectral imaging have received considerable attention for harvest date anticipation based on berry ripeness assessment (e.g. [Navrátil and Buschmann, 2016; Fernández-Novales et al., 2019; Kalopesa et al., 2023]). The major interest of these methods is that they eliminate the need for solute extraction and physico-chemical tests, and make it possible to objectivise the heterogeneity of maturities at plot level. However, such data acquisition may be practically as tedious as harvesting representative samples. It also misses the kinetics of volume growth, which is critically needed to predict yield and distinguish the sugar accumulation phase from its final concentration. Alternatively, berry volumes, as well as their colour, can be deduced from an image of a grapevine bunch. Since these morphological features are linked to berry ripeness, ripening dynamics could be measured by analysing time-series of such images. While the efficiency of this non-destructive approach was demonstrated with manual annotation of the images [Lou et al., 2016], only the automation of such tasks would allow a large enough sampling to get a representative view of the ripening process and its variability.

The first task to be automated is the detection and segmentation of individual berries. This task is challenging, due to the natural variability of the aspect of berries (e.g. shape, size, colour, degree of light exposure) and to the fact that they frequently overlap with other berries and plant parts. Deep-learning has shown to be an effective solution to this problem for a number of fruits such as oranges [Ganesh et al., 2019], blueberries [Gonzalez et al., 2019; Ni et al., 2021], apples [Jia et al., 2020; Gené-Mola et al., 2020], strawberries [Perez-Borrero et al., 2021] and grapevine berries [Shen et al., 2022]. In all these studies, an instance segmentation model (e.g. Mask R-CNN [He et al., 2017]) was trained on manual annotations of visible fruit parts to retrieve the apparent contour of each fruit. This strategy is suitable for measuring their colour [Shen et al., 2022], counting them to estimate yield [Zabawa et al., 2020], or locating them for automatic fruit picking [Tang et al., 2020]. However, it misses the occluded parts of berries that are partially covered by neighbouring fruits, which frequently occurs in ordinary bunches, thus preventing the deduction of statistics related to their real shape such as volume. To cope with this, [Miao et al., 2021] used ellipse fitting as a post-processing of the segmented contours to infer a plausible intrinsic contour of individual berries. Alternatively, deep-learning models can be trained on annotations guessing the shape each fruit would have in the absence of occlusions, so that predictions of the segmentation model directly infer complete fruit shapes, including their hidden parts [Bargoti and Underwood 2017; Liu et al., 2018; Dolata et al., 2021]. The annotation protocol and the extent to which the hidden parts can be deduced from the visible ones are crucial in such cases, as annotation errors will be learned by the models and will directly alter predictions. Higher level of occlusions can be addressed by training the model with synthetic images for which various levels of occlusion can be generated, by artificially superposing images of isolated fruit and other plant elements [Hondo et al., 2022] or by rendering plant models in a 3D graphics software [Barth et al., 2018].

The second task to be automated is the tracking of segmented berries over successive time steps, to deduce individual volume and colour kinetics. The majority of fruit tracking algorithms addressed the issue of matching segmented instances between different viewpoints, or over time on short videos (seconds to minutes) of several frames per second [Liu et al., 2018; Wang et al., 2019; Zhang et al., 2022]. [Hondo et al., 2022] managed to track apples over periods of several weeks, but for a very limited number of instances (two well separated apples), which is far from the issue of tracking dozens to hundreds of overlapping instances over a long period of time, as needed for following berry ripening.

In this paper, we introduce a fully automated method to measure and track the size and colour of individual berries on time-lapse images of grapevine bunches. The method starts with a detection model to recognize berries that are sufficiently visible to reasonably infer their size. Second, a segmentation model was trained to infer both the visible and hidden contours of individual berries, using a training dataset derived from a fast and original annotation method. Ellipses are further fitted on the segmented contours to compute position and shape parameters for each berry. Finally, we adapted a tracking algorithm to assign time-consistent labels to the detected berries while handling global deformations of the bunch. This method was tested on image time-series acquired at the PhenoArch platform [Cabrera-Bosquet et al., 2016], to assess the quality and the limits of the method at quantifying individual berry growth kinetics. We finally showed how this unprecedented data analysis can provide new insights on the ripening dynamics of grape berries.

## Materials and methods

### Plant material, image acquisition and dataset composition

The complete pipeline (segmentation and tracking) was tested on an image dataset from two independent experiments conducted in 2020 and 2021, spanning 51 and 32 days respectively, each containing 9 grapevine (*Vitis vinifera* L.) plants. For each plant, images of a selected grapevine bunch were taken every 8 hours. An additional dataset including bunch images of 78 grapevine genotypes from a diversity panel maximising genetic diversity [Nicolas et al., 2016] was used to robustify the training and evaluation of the berry segmentation pipeline (without tracking). All experiments were conducted in the PhenoArch phenotyping platform (https://www6.montpellier.inrae.fr/lepse_eng/Phenotyping-platforms-M3P/Montpellier-Plant-Phenotyping-Platforms-M3P/PhenoArch), hosted at the M3P (Montpellier Plant Phenotyping Platforms) [Cabrera-Bosquet et al., 2016].

For each plant shot, a set of 12 RGB images (2048×2448 px) were taken around the grapevine bunch with a regular 30° rotational offset, using an imaging cabin of PhenoArch. The cabin involves an RGB camera (Grasshopper3, Point Grey Research, Richmond, BC, mounted in a robotized XYZ arm and LED illumination (5050–6500 K colour temperature). Images were captured while the plant was rotating at constant rate (20 rpm) using a brushless motor (Rexroth, Germany). For each plant, the bunch position was manually recorded at the beginning of the experiment, and a robotic arm (see [Brichet et al., 2017] for details) was then used to automatically position the camera to a fixed time-consistent position along the experiment allowing to get a detailed shot of the bunch.

### Detection, segmentation and features extraction of individual berries

The objective of this step is to i) detect berries suitable for shape inference, defined as berries with more than half of their contours visible in a grapevine bunch image, ii) infer their complete shape, and iii) extract features that allow quantifying their size and colour. The first two sub-steps rely on deep-learning models which have to be trained on annotations of complete berry contours inferring their hidden part.

#### a) Construction of the annotation dataset

The annotated dataset contains 159 images, sampled from the three experiments (Table 1). The sampling was done to best cover all stages of growth, and maximise the visual diversity of the berries in the dataset in terms of size, shape, colour, texture, blurring and shading. It also includes various levels of occlusions between berries or with other plant organs and objects.

**Table 1-.** Annotated dataset

| experiment<br>year | n. of images | n. of plants | n. of genotypes | n. of annotated<br>berries |
| --- | --- | --- | --- | --- |
| 2020 | 8 | 8 | 1 | 536 |
| 2021 | 33 | 22 | 3 | 1670 |
| 2022 | 118 | 109 | 78 | 3928 |

A total of 6,134 berries were manually annotated as polygons using Labelme [Wada 2018]. Similarly to [Miao et al., 2021], only berries with at least half of their contours visible were annotated. Berries that did not reach pea size stage were rejected based on the assessment of their morphological characteristics, as they are not relevant for studying ripening. For each berry, an average of only 8 points (at least 5) were placed along the uncovered parts of its contours (Fig. 1A). Then, least-square ellipse fitting [Fitzgibbon et al., 1999] was used to fit 5 ellipse parameters (*x*_*e*_, *y*_*e*_, *w*_*e*_, *w*_*e*_, *a*_*e*_) to the set of points (Fig. 1B; blue lines), with (*x*_*e*_, *y*_*e*_) the centre coordinates of the ellipse, *w*_*e*_ and *h*_*e*_ the respective length of minor and major ellipse axis, and *a*_*e*_ the ellipse rotation. *w*_*b*_ and *h*_*b*_ were further deduced as the width and height of the smallest box enclosing the ellipse.

**Fig. 1.**
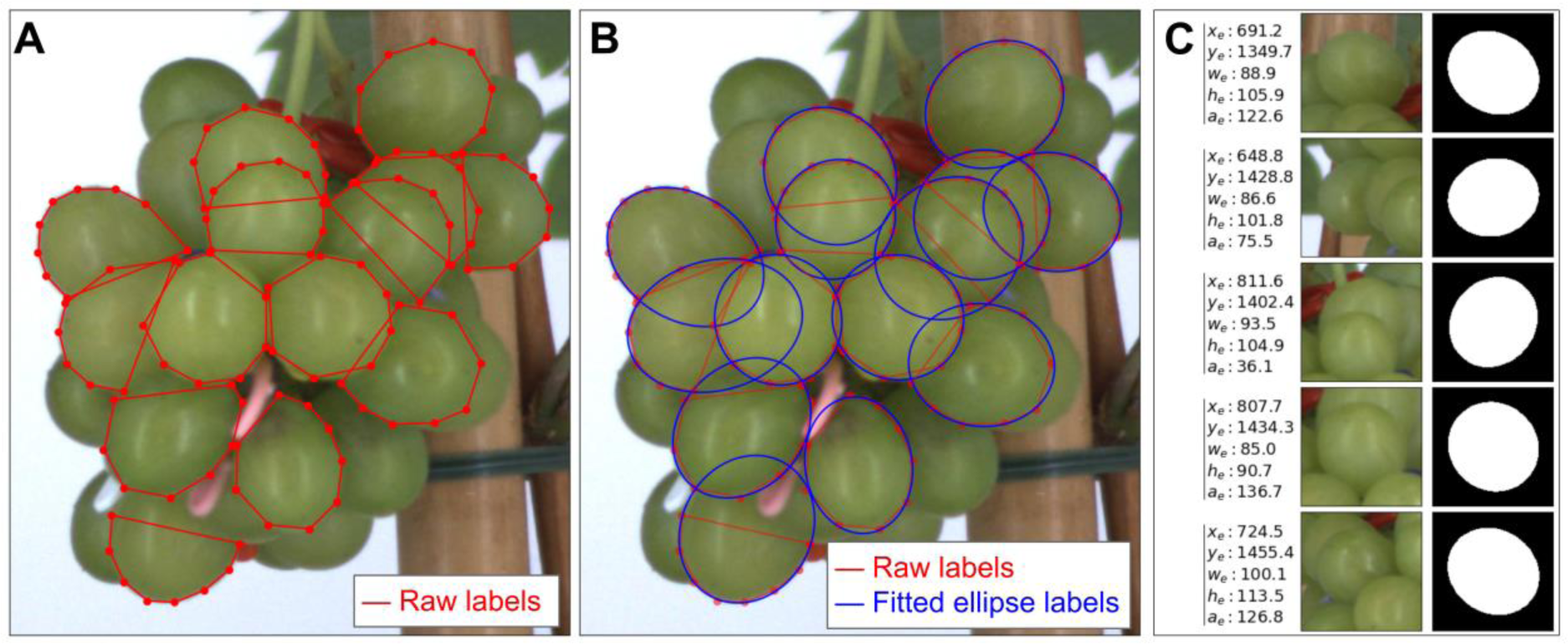
Berry annotation procedure. **A** Raw labels, consisting of simple polygons (5 to 10 points) drawn manually along the edges of berries with at least 50% of their contour visible. **B** Guessed actual contour of berries, obtained by an automatic ellipse fitting (blue) on the annotated points. **C** Instances generated from the annotations dataset, used to train the segmentation model. Each instance corresponds to one berry, for which we show the fitted ellipse parameters, the image input and the targeted binary segmentation mask.

This dataset (Table 1) was then split into training (129 images; 4,447 labels), validation (10 images; 814 labels) and test (20 images; 873 labels) subsets. Each subset includes different plants, to better assess the generalisability of the detection and segmentation models.

#### b) Detection of measurable berries

To detect measurable berries on an image of a grapevine bunch, a Yolov4 deep-learning object detection model [Bochkovskiy et al., 2020] was trained to find bounding boxes around berries with at least 50% visible contour in 416×416 px sub-parts of the image (Fig. 2A-B).

**Fig. 2.**
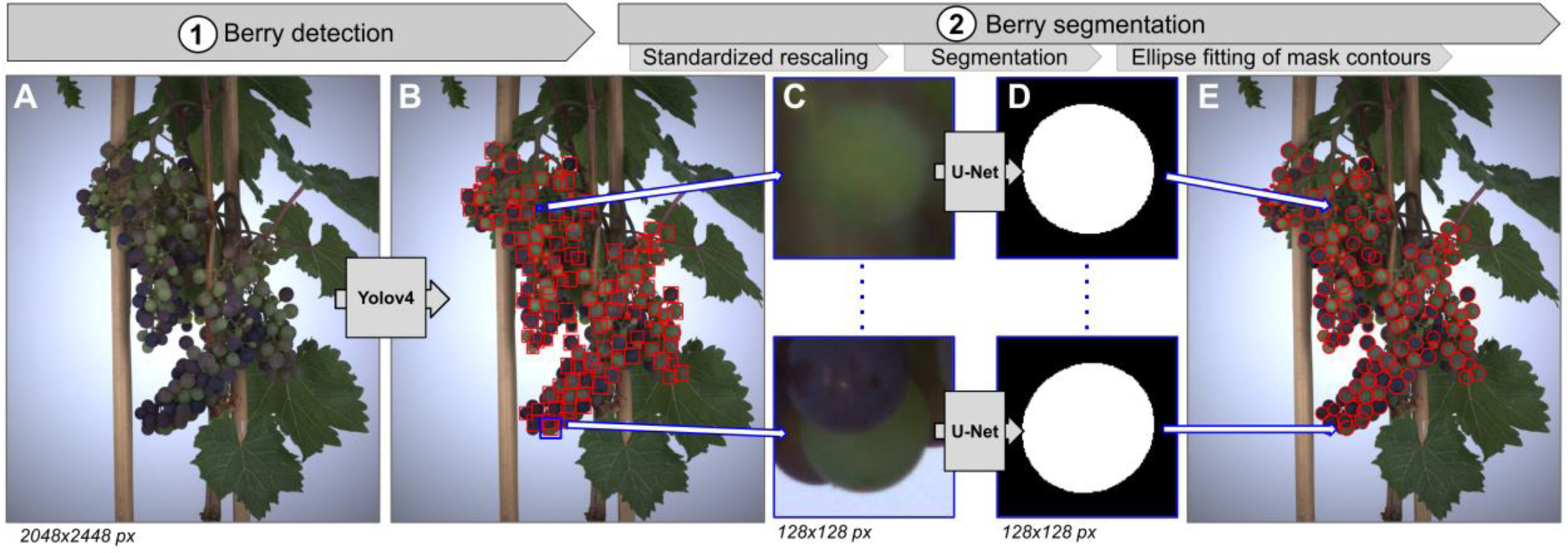
Berry detection and segmentation pipeline. **A** RGB image of a grapevine bunch acquired in the PhenoArch platform [Cabrera-Bosquet et al., 2016]. **B** Bounding boxes (red rectangles) detected by a Yolov4 deep-learning model trained to identify berries with at least 50% visible contour. **C** Vignettes cropped around the centre coordinates of detected boxes, and resized to 128×128 px. The resizing ensures that berries occupy a similar space in the vignette regardless of their size. **D** Binary segmentation masks predicted by a U-Net deep-learning model on berry vignettes. The model was trained to infer the shape of berries in the absence of occlusions. **E** Ellipse fitting of the contour points extracted from a segmentation mask, and projection of the ellipse (red) on the original image.

20,000 training instances were generated by cropping 416×416 px sub-parts of the training images, each being labelled by the list of parameters of the boxes entirely included in it. This dataset was further augmented with random adjustments of vignettes hue, saturation and brightness, and random flips of image-label pairs. It was then used to train the model, using the yolov4-tiny architecture and default hyperparameters [Bochkovskiy et al., 2020]. Model weights were stored every 500 iterations for a total of 65,000 iterations. The weights leading to the highest Average Precision (AP) on the validation dataset were saved.

For predictions, the 2048×2448 px source image is split into image sub-parts cropped over the entire pixel range with a maximum spacing of 270 px, which are then fed to the detection model, resulting in a set of predicted parameters of the box dimensions 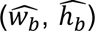 and centre coordinates 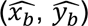. Because of sub-image overlaps, the same berry can be detected more than once. To remove these redundancies, non-maximum suppression is used to avoid having box pairs with an intersection over union above 70%. Berries detected with a confidence score below a threshold *s* = 0.89 are filtered out. This value of *s* was chosen to maximise the F1-score on the validation subset.

#### c) Segmentation of berries

For each detected berry, a square vignette of size 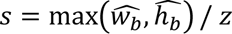 is cropped around its box centre coordinate 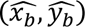. A constant value *z* = 0.75 is used to ensure that all berries are entirely contained within their respective vignette, and occupy a similar space regardless of their size (Fig. 2C). Each vignette is then resized to 128×128 px by bilinear interpolation, and fed to a U-Net [Ronneberger et al., 2015] deep-learning model with a VGG16 [Simonyan and Zisserman 2015] backbone. The model was trained to output a binary mask representing the shape of the berry as if it were not occluded by any other element present in the image (Fig. 2D).

To train the segmentation model, 40,000 vignettes were extracted from the annotation dataset using the cropping method described above. Elliptic mask labels were directly generated using the annotated ellipse parameters (Fig. 1C). Random noise was applied to the centre coordinate and value of *z* during cropping, to help the model handle detection inaccuracies. This was supplemented by the augmentation scheme explained earlier in the detection section. Therefore, all the masks generated had ellipse shapes of similar sizes, in order to restrict the learning domain of the model. These vignettes and mask labels were then used as inputs and output to train the model, using categorical cross-entropy loss, Adam optimizer, and a learning-rate of 0.0001. The number of iterations was automatically chosen with early stopping, and the model weights leading to the minimal validation loss were saved.

#### d) Extraction of berry morphology and colour features

Assuming that the resulting mask has an elliptic shape, its contour points are extracted as in [Suzuki and Be 1985], to fit 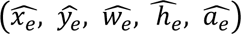 ellipse parameters [Fitzgibbon et al., 1999]. These parameters are then rescaled to the original image coordinate space (Fig. 2E). For each berry the following features are computed:

**Colour:** the raw hue *h*_*raw*_ of a berry is computed as the circular mean of the hue angle of the pixels contained inside the ellipse, after removing the pixels that are less than *dp* = max(3, *w*_*e*_/4) px away from the ellipse’s edges, and removing the pixels shared by other ellipses. Given *h*_50_ = 100° the mean value of *h*_*raw*_ for grape berries that are halfway through their colour change from green to black in our dataset, the centred berry hue *H* is defined as:

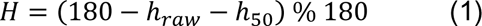

**Volume:** Berry volume *V* is estimated as the volume of the sphere that has the same projection area *A* as the ellipse fitting the individual berry shape, as in [Dubois et al., 2022]:

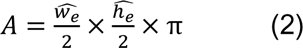

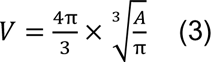

Assuming that the distance between a bunch and the camera was uniform across plants, and relatively large compared to the differences in distance to the camera across berries, a constant calibration factor of 3.94 × 10^−6^*mL*. *px*^−3^ was used to express *V* in mL.

### Time-lapse tracking of individual berries

This step aims to track individual berries over successive segmented images of a grapevine bunch, that is to associate a unique label to each berry over time (Fig. 3A, 3E). To that end, three independent methods were combined (i.e. Baseline, Registration and Matching Tree). First, an original algorithm (Matching tree) was used to both find the best starting point *t*_*root*_ to initialise the labels, and optimally reorder the way these labels are propagated to other time-steps (Fig. 3D). This algorithm is based on the construction of a distance matrix that quantifies the dissimilarity between all possible pairs of time steps (Fig. 3C). The tracking itself is based on an iterative matching of the central coordinates of the ellipses between two time steps (Baseline), and includes a pre-processing step to better manage the global movements of the bunch (Registration, Fig. 3B).

**Fig. 3.**
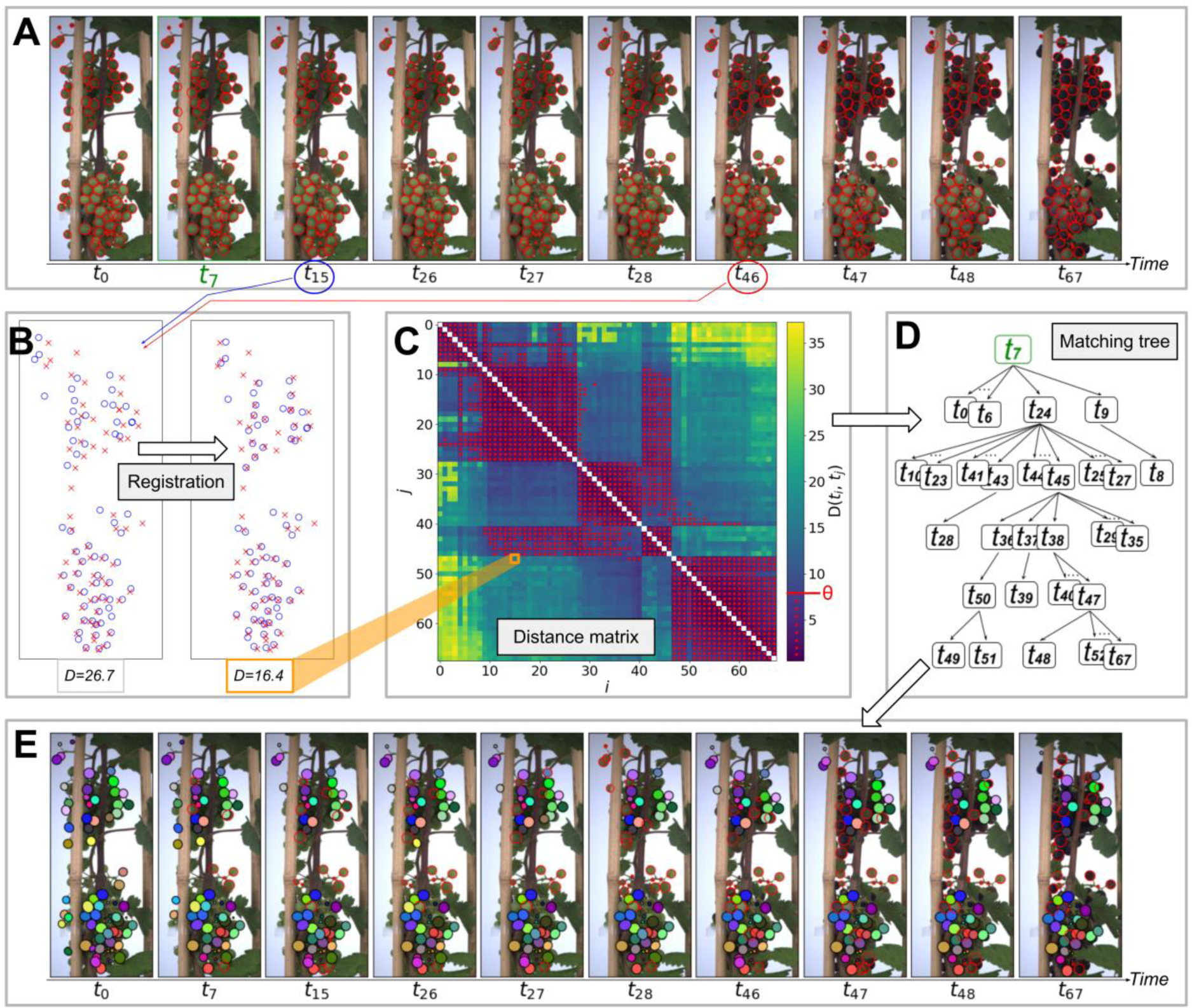
Berry time-lapse tracking pipeline. **A** 10 segmented RGB images sampled from a 68 images time series representing the evolution of one bunch over time. Raw images were captured with a median interval of 8h. **B** Scatter plots of the coordinates of the berry ellipses centres detected at two time steps (*t*_15_; blue circles, *t*_46_; red crosses), before (left) and after (right) registration. The distance metrics *D* between the two point sets is given below each plot. **C** Heat map of the distance matrix, storing the distance between all pairs of time-points after registration. Red points correspond to matrix values below the threshold θ = 8*px*. **D** Matching tree, determining the order in which labels are propagated during tracking. Each rectangle represents a time-step. The highest one corresponds to *t*_*root*_, used to initialise tracking labels. **E** Labelled segmented images after tracking. Each colour corresponds to one tracking label. Segmented berries without label (no match found with *t*_*root*_) are drawn as red empty ellipses.

#### *a) Baseline:* matchings of berry centre coordinates between two time steps

For any time-point *t*, the segmentation provides a point set *S*_*t*_ = {*c*_*k*_} containing the ellipse centre-point coordinates 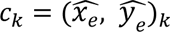 of each berry detected. Assuming all berries remain in the image frame with approximately constant relative positions, the tracking of berries from time steps *i* to *j* is treated as a bipartite matching of point sets *S*_*i*_ and *S*_*j*_. The correspondence between two point sets is done by associating to each centre *c*_*i*_ in *S*_*i*_ its nearest neighbour *c*_*j*_ in *S*_*j*_ in euclidean distance.

Each point can only be paired once, the closest pairs are matched first, and pairs with a distance above a threshold δ = 16*px* are discarded. The value of δ was chosen as the quarter of the median value of *w*_*e*_ in our annotation dataset, with the idea that such a low value strongly limits mismatches, even in dense areas of the bunch. This algorithm can be applied successively to pairs of sets (*S*_*t*_, *S*_*t*+1_) along a time-series of *N* images, to propagate the correspondence of the initial set of labels.

#### *b) Registration:* estimating global bunch deformations prior to matching

Even if the berries in a bunch maintain the same relative arrangement, their absolute positions may change between two time steps *i* and *j* due to relative movements of the bunch and the camera, or due to internal deformations and movements of the bunch. Assuming that the resulting deformation of the point cloud in the image coordinate system is affine, the Coherent-Point Drift algorithm [Myronenko and Song, 2010] was used to realign the two sets prior to matching, by finding the affine transformation 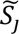 of *S*_*j*_ that minimises the distance to *S*_*i*_ (Fig. 3B).

#### *c) Matching tree:* processing the time steps in an optimal order

The matching algorithm can be applied to any pair of sets (*S*_*i*_, *S*_*j*_) from the time-series. Propagating the matching to successive pairs (*S*_*t*_, *S*_*t*+1_) in chronological order is a common choice in multiple object tracking [Luo et al., 2021]. However, this option may not always be optimal, as in our case where the camera may for example move unexpectedly at a time step and then return to its original position (see video in Additional File 1 for examples). Here we propose to match the most similar pairs of sets in priority, to avoid errors that could occur and propagate from a pair of sets that are too dissimilar. To do so, we define a metric *d* quantifying the distance from *S*_*i*_ to *S*_*j*_, based on the euclidean distance function *e*:

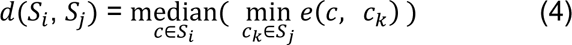

Since *d* is not commutative (e.g. one point in a set could be the closest neighbour of many points in the other set), we define the distance *D* between two sets *S*_*i*_ and *S*_*j*_ as the average of *d*(*S*_*i*_, *S*_*j*_) and *d*(*S*_*j*_, *S*_*i*_). Unlike during the matching *(Baseline)*, the computation of *D* does not involve a bipartite pairing of points, which is time consuming because of the iterative nature of the algorithm. It therefore allows for a fast quantification of the dissimilarity 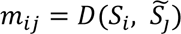 between all pairs of time-steps, which are stored in a distance matrix *M* = (*m*_*ij*_) (Fig. 3C). This matrix is used to arrange the order of the successive matchings through a layered tree (Fig. 3D). Unlike a linear ordering of the successive time-steps, arranging them through a tree structure reduces the average number of intermediate steps between *t*_*root*_ and other time-steps, thus limiting the risk of propagation of matching errors.

The matching tree contains a root node *S*_*root*_, and each node has a depth *k* equal to the length of its path to the root. To select the nodes of depth *k*, we iteratively connect the candidate set *S*_*i*_ to another set *S*_*j*_ of depth less than *k*, such that *d*_*min*_ = min(*M*_*ij*_, *M*_*ji*_) is minimised. This is repeated as long as *d*_*min*_ < θ, with θ = 8*px* a threshold controlling the ratio between the width and depth of the tree. If no candidate set meets this criterion for depth *k*, a single *long-distance* edge is built between layers *k* − 1 and *k* with the minimum possible distance. This process is iterated for successive depths until the tree contains all sets of *S*. *t*_*root*_ is selected exhaustively as the value allowing to place the most nodes in the tree before reaching a *long-distance* edge, and secondarily by maximising the number of points in *S*_*root*_.

### Evaluation of the method

To evaluate the berry detection on a given image, the predicted ellipses whose intersection over union are greater than 0.5 with a labelled ellipse are classified as True Positives (TP). The remaining predicted and labelled ellipses are respectively classified as False Positives (FP) and False Negatives (FN). Precision, Recall and F1-score metrics are then deduced as follow:

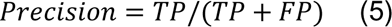

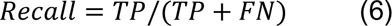

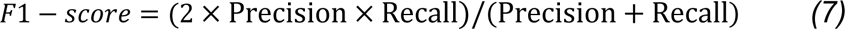

For segmentation evaluation, the area of the segmented ellipses was compared with ground-truth observations using the following metrics: bias, root mean-square error (RMSE), mean absolute percentage error (MAPE) and coefficient of determination (*R*²).

Berry tracking was evaluated by two metrics, namely the coverage *T*_*c*_ and the precision *T*_*p*_. *T*_*c*_ is defined as the percentage, over the full time-series, of segmented berries that could be matched by the tracking algorithm to a berry segmented at *t*_*root*_ (coloured ellipses in Fig. 3E). *T*_*p*_ is the percentage of labels that point to the same berry over time, and was estimated by manually checking random samples of 10 time-steps per bunch in each time-series. Both metrics were computed using, for each bunch, the time-series from the camera angle providing the largest number of segmented berries.

One bunch of the 2020 experiment was further analysed to assess the potential of the method at capturing and quantifying berry development and its asynchrony. A total of 81 berries were extracted, combining three camera angles (120° interval), and after selecting berries tracked over at least 90% of the experiment duration. For each berry, an 8-days moving median was used to smooth the raw volume measurements over time (Fig. 7A and Additional File 2A; red curves), and a MAPE value was computed between the raw and smoothed volume data. The 10% berries with the highest MAPE were excluded from the analysis to reduce the noise, resulting in a final dataset of 73 berries. For each kinetics of a variable *X* (either *V* or *H*), a relative kinetics *X*_*r*_ and a scaled kinetics *X*_*s*_ are computed as:

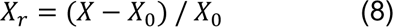

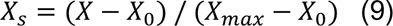

*X*_0_ is the median of *X* during the first 8 days. *X*_*max*_ is the maximum of the smooth kinetics for

*V*, and the median of the last 8 days for *H*. All the method conception and the data analysis were performed in Python.

## Results

### Deep-learning segmentation allows accurate and robust shape inference of partly hidden berries

Berry segmentation was performed on the 2020 and 2021 datasets (21,744 images), resulting in an average detection of 64 berries per image. Fig. 4 provides some examples of detection on the test subset, showing that the model was able to infer the full contour of overlapping berries from different genotypes varying in size, shape and aspect, even when these contours were not fully visible. Predictions on the full test subset were compared with ground-truth annotations to quantify both detection and segmentation accuracies.

**Fig. 4.**
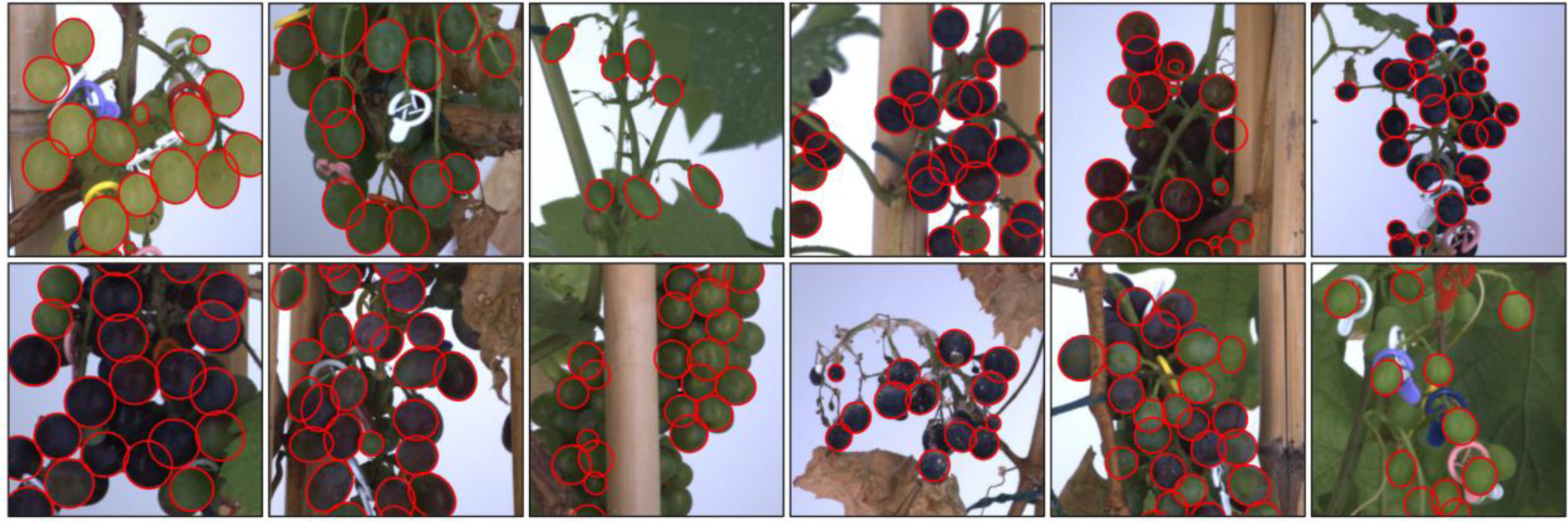
Examples of segmented grapevine bunch images. Output of the berry detection and segmentation pipeline on bunch images from 12 grapevine cultivars. Images come from the test subset, and none of these cultivars were used to train the model. Only a 500×500 px subpart of each image is shown.

The detection of measurable berries had a Precision of 92.3% and a Recall of 87.5% on the test subset, resulting in a F1-score of 89.8%. The remaining errors (64 FPs, 109 FNs) were further investigated (e.g. Fig. 5) through a manual classification (Additional File 3A). This revealed that most errors (59% of FPs and 52% of FNs) correspond either to berries with a visible contour fraction within a 50±10% range, or to small underdeveloped berries (around pea size stage). Both situations are close to the selection criteria used when annotating berries, and the assessment of whether or not these criteria have been crossed may be ambiguous for both the annotator and the model. For FPs (i.e. detected but not annotated berries), errors were evenly distributed across berry sizes (Additional File 3B). 56% of them correspond to berries within the 50±10% visible contours range, sometimes due to an error by the annotator detected a posteriori. Considering that berries within this 10% error range are still good candidates to shape inference, the precision of the method at detecting measurable (even if not annotated) berries can thus be re-estimated to 96.0% (F1-score = 91.8%). Concerning FNs (i.e. missed detections), pea sized berries alone account for 27% of the cases, which result in a slight under-representation of this class in the histogram of the size of berries detected. (Additional File 3B).

**Fig. 5.**
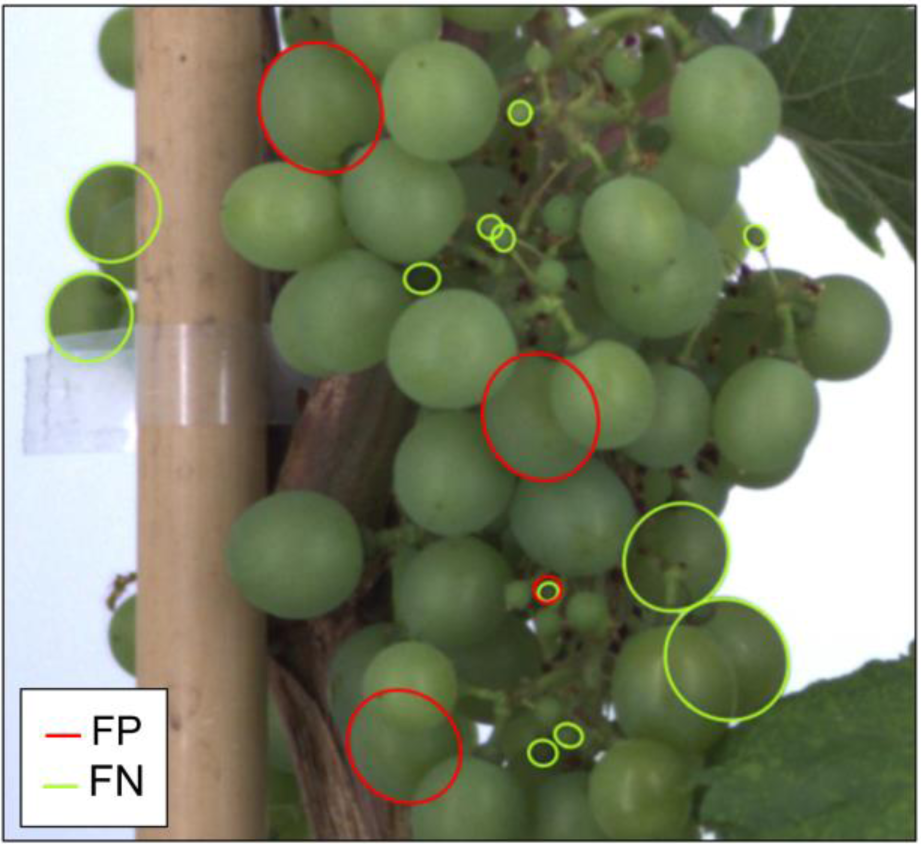
Example of mismatches between the berry detections and annotations. False positives (FP, red) and false negatives (FN, green) found when comparing berries detected by the pipeline to manually annotated berries, on a grapevine bunch image from the test subset. Only a subpart of the full image is shown.

The area of the ellipses segmented by the model closely matched those of the manual annotations on the test subset (Fig. 6; MAPE=4.1%, *R*²=0.976), with a low bias of −32px². This demonstrates that the segmentation model was able to accurately infer the size of berries with up to 50% of their contours hidden. A similar MAPE around 4% was obtained on genotypes either present (n=440) or absent (n=363) from the training subset, suggesting that the segmentation generalised well to the genetic diversity in our dataset.

**Fig. 6.**
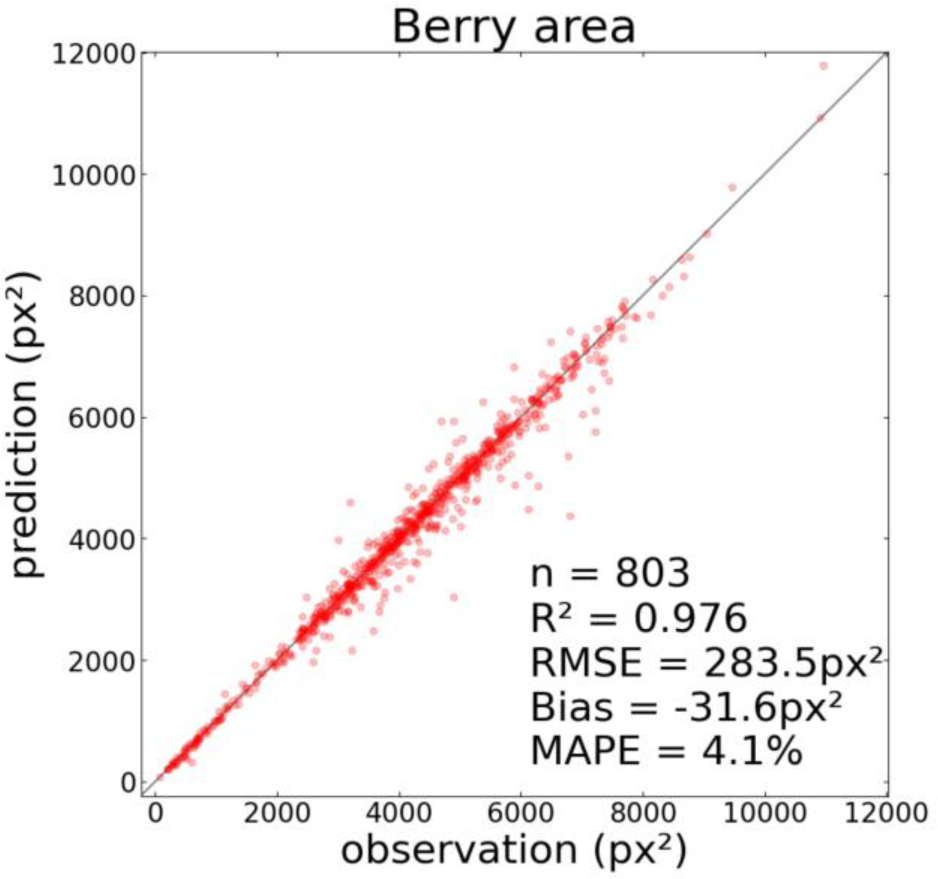
Accuracy of berry area measurement. Comparison of the area of manually annotated berries (observation) with those from the detection and segmentation pipeline (prediction). n = number of points, RMSE = Root-Mean Square Error, MAPE = Mean Absolute Percentage Error.

### An almost error-free tracking of 50 to 80 % of the segmented berries

Berry tracking was performed on 9 grapevine bunches from different plants in each of the 2020 and 2021 experiments, observed over an average of 65 and 136 time-steps respectively, with the same median interval of 8h between images. The video from Additional File 1 shows examples of tracking outputs for three different bunches. It was observed that image time-series exhibit periods during which the relative position of the camera and the plant remain stable, resulting in a fixed positioning of the bunch in the image, but also include irregular movement of the plant and of the camera despite the robotisation of the image acquisition, combined with irregular movements of both the bunch and the leaves.

The coverage *T*_*c*_ of the berry tracking method was assessed for each of the 18 bunches (M4, Table 2). The individual effects of the method components *(Registration, Matching tree)* were evaluated by re-running the tracking without them (M1 to M3, Table 2). Two subsampling scenarios were also used to assess the effect of increasing the time step from 8 to 80h (S1, Table 2) and restricting the time-series to periods of stable shooting conditions (S2, Table 2). These periods were manually identified by careful examination of the stability of the image acquisition over time.

**Table 2-.** Coverage index (*T*_*c*_) of the tracking method (M4) for two experiments. Lines correspond to different combinations of the tracking algorithm elements (M1 to M3) and two sub-sampling strategies of the data (S1, S2).

| Tracking method | Mean coverage ($T_c$) | |
| --- | --- | --- |
| | 2020 ($n=9$) | 2021 ($n=9$) |
| M1: Baseline | 5.1% | 40.8% |
| M2: M1 + Registration | 32.3% | 62.8% |
| M3: M1 + Matching tree | 35.6% | 63.2% |
| **M4: M1 + Matching tree + Registration** | **53.4%** | **74.2%** |
| S1: M4 on an increased time-step (8 to 80h) | 54.1 % | 70.4% |
| S2: M4 (and M1) restricted to k stable periods<br>($\bar{k}$ : mean value of k) | 77.1% (24.1%)<br>($\bar{k}=3.4$) | 82.0% (52.2%)<br>($\bar{k}=1.8$) |

*T*_*c*_ had an average value of 53.4% and 74.2% for 2020 and 2021 experiments respectively. The precision was very high in both 2020 (*T*_*p*_=96.7%, 623 labels) and 2021 (*T*_*p*_=99.2%, 793 labels) experiments. This indicates that the tracking method is more accurate than exhaustive, which is appropriate for studying berry growth kinetics, since an accurate monitoring of a representative subsample of berries is sufficient to reflect the whole bunch dynamics. Such a high precision might be ensured by the low value chosen for the distance threshold δ which determines whether two segmented berries can be matched. Using point-set registration (Fig. 3B) and a matching tree (Fig. 3D) during tracking both contribute to maintain sufficiently high coverage, as it increases *T*_*c*_ by a factor of 10.4 and 1.8 for 2020 and 2021 experiments respectively, compared to a regular succession of point-sets matchings (Table2; M2-4 vs M1).

However, a significant amount of segmented berries (2020: 46.6%, 2021: 24.8%) remained unmatched to *t*_*root*_. The time interval between images was not likely to explain these losses, since a 10-fold decrease in the image frequency did not significantly modify *T*_*c*_ for the same duration (Table2; M5). Instead, further examination of the distance matrices computed during tracking highlighted periods with a strong temporal consistency (i.e. low distance between point sets of ellipse centres), separated by abrupt transitions which were often associated with a drop in *T*_*c*_ (Additional File 4). 30 transitions were empirically annotated using these matrices, to identify their cause on their corresponding images (Additional File 4A; red lines). Most transitions coincided with a bunch rotation (70%), a strong shift in camera position causing berry apparitions or disappearances (13%), or a deformation within the bunch (10%). These situations correspond to the actual limitations of our registration method, but most of them could have been avoided by a better management of the experimental conditions. Performing tracking independently in each time consistent period increased *T*_*c*_ to 77.1% and 82.0% for 2020 and 2021 experiments respectively (Table2; M6). These metrics probably reflect the performance of our method under experimental conditions where the instability of the image acquisition is better managed.

### Robust measurement of single berry dynamics, differing from the usual “mean berry” approach

Combining the tracking labels with the features extracted from the segmented berries allowed to monitor the growth of a single berry over time with high accuracy and temporal resolution, both in terms of volume and colour (Fig. 7). While volume measurements can be noisier due to variations of just a few pixels in the image, colour measurements are more reliable because they are derived from averaging a larger number of pixels. These kinetics exhibit smooth patterns over time, using high frequency measurements of a large number of berries in several bunches (Additional File 2), which supports the suitability of this method to high-throughput phenotyping conditions.

**Fig. 7.**
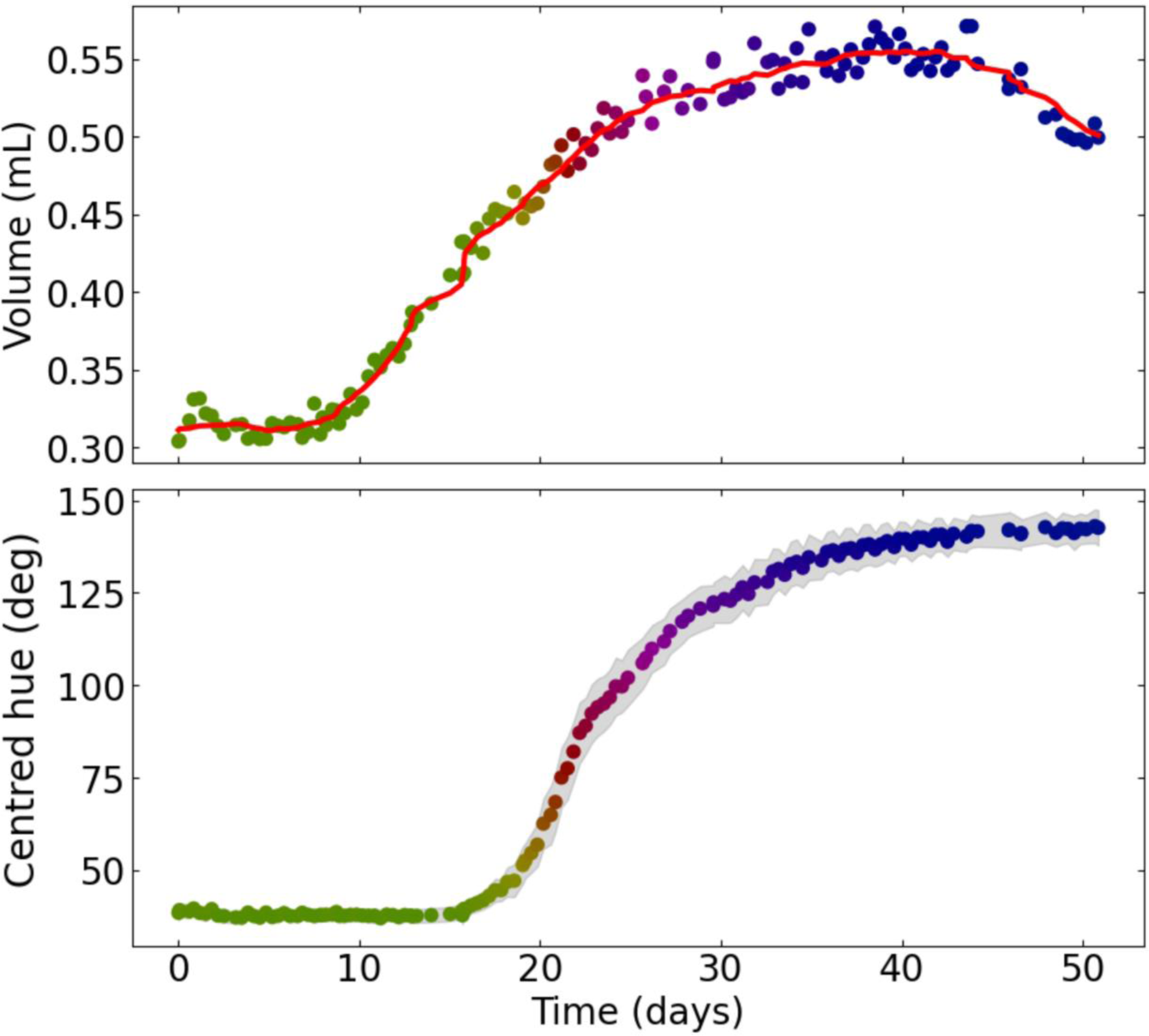
Growth and coloration kinetics of an individual grapevine berry. Volume (**A**) and Centred hue (**B**) measured over time on an individual berry, after running the full image analysis pipeline on a time-series of 138 bunch images. All points are coloured using the corresponding average hue values. In **A**, the red curve corresponds to an 8-days moving median smoothing. In **B**, the grey area corresponds to the standard deviation of the centred hue value observed within the berry segmentation mask.

The potential of the method to reveal new characteristics of berry ripening and bunch population structure was further assessed on 73 individual berries tracked on a bunch from the 2020 experiment, on the *Vitis vinifera* cv Alexandroouli, a black hermaphrodite cultivar of Georgian origin used for wine making (https://www.vivc.de/?r=passport%2Fview&id=263). The bunch was observed for a period of 50 days, corresponding to the second growth period of berries, which goes from the end of the green phase to over-ripening (Fig. 8-9). To our knowledge, this is the first report on the growth and colouring dynamics of a statistically relevant number of individual fruits. We confirm here that, at similar developmental stages, the individual berry volume exhibits 2-3 fold variations with considerable deviations from gaussian distribution inside a single bunch (Fig. 8A). On average in this bunch, an individual berry increased its volume by 60% (Fig. 8A) in 18 days before reaching its maximal volume (Fig. 8C; grey dotted line), and underwent a more or less intense shrivelling period once phloem unloading in berries definitively stopped [Savoi et al., 2021]. Such a relative expansion rate is in line with the approximate doubling of berry volume during the ripening of most *V. vinifera* cultivars [Houel et al., 2013; Bigard et al., 2019; Bigard et al., 2022], which took three weeks to complete on individual fruits of Meunier, Syrah or ML1 [Shahood et al., 2020], Cabernet Sauvignon and Pinot [Friend et al., 2009]. Further studies are needed to establish if the slightly shorter growth duration and expansion of Alexandroouli’s berry is truely of genetic origin, or is the result from tests on fairly young own rooted potted plants in greenhouse conditions. In any case, we confirm here that the typical duration of the second growth period of an individual berry lies within 30-50% at best of the consensual duration of ripening in textbooks, which is routinely inferred upon averaging hundreds random berries representing fruit diversity at the parcel scale (i.e. [Krasnow et al., 2013; Suter et al., 2021; Fasoli et al., 2018]). Present data even shows that for a single bunch, which undoubtedly underestimates asynchrony at plot scale, the global growth curve recalculated for all detected berries noticeably overestimates the average duration of the second growth period (Fig. 8C; red star) and underestimates the maximum growth rate (Fig. 8D; red star). These statistical biases clearly result from adding the asynchrony to the real, but previously unknown, duration of the second growth period on average representative samples. Moreover, the fact that asynchrony and growth duration last approximately as long in a single bunch (Similar ranges for the y-axis of Fig. 8B-C) means that berries at the same phenological stage (e.g. all growing) cannot be collected, which is a major drawback of conventional samples for tackling all aspects of fruit development biology. Real time monitoring of berry growth allows to constitute synchronised berry samples more conveniently than marking each berry fecundation or softening dates. In this respect, growth resumption precedes coloration by more than four days on average, but the delay can vary from one day up to two weeks (Fig. 9). This variability clearly limits the use of coloration as a proxy for the berry internal clock [Vondras et al., 2016]. Moreover, present original insights on the dynamic structure of berry cohorts within a bunch allow us to test if the ripening program accelerates in the late berries [Gouthu et al., 2014], or if ripening berries compete for water or photoassimilates. Such possibilities should be rejected here, as berry relative expansion didn’t diminish with the timing of growth resumption, nor with the number of growing berries collectively acting as a possible competitive sink at this date (Fig. 8B; R²=0.02). Nevertheless, individual berries largely differed in their maximum relative expansion, which was clearly influenced by their maximal growth rate (Fig. 8D; R²=0.59), not by growth duration (Fig. 8C; R²=0.002). This first approach on the dynamic structure of berry population based on the discretisation of single berry dynamics clearly constitutes a paradigm shift from modelling the future crop as an average ideal fruit [Zhu et al., 2019].

**Fig. 8-.**
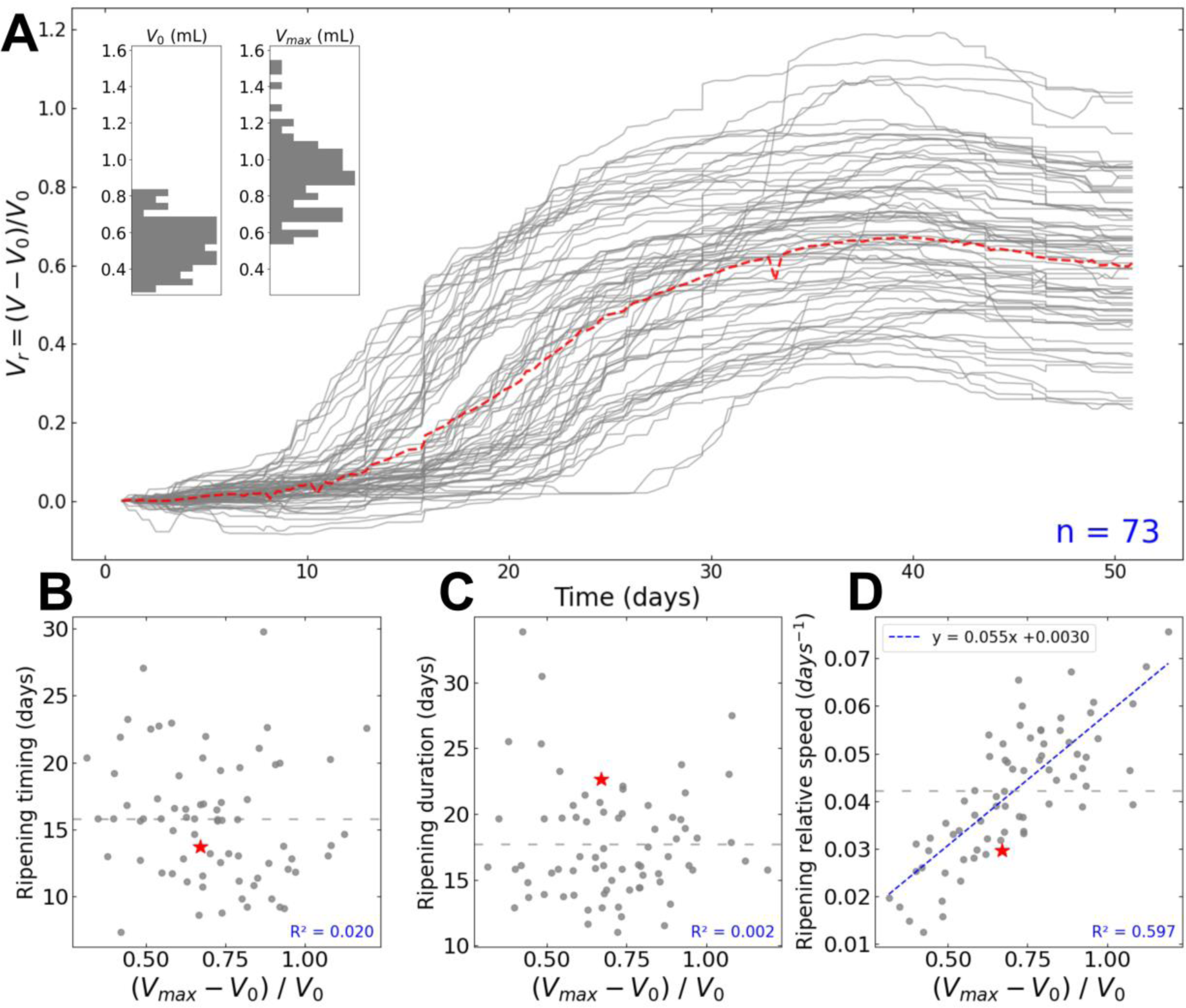
Individual growth kinetics of berries within a grapevine bunch. Smoothed volume (*V*) kinetics were automatically computed for *n* = 73 berries, by applying the full image analysis pipeline on time-series of 138 images from 3 different camera views (120° difference) of the same grapevine bunch. The *ripening timing* corresponds to *t* = *t*(*V* = 0.15). The *ripening duration* is extrapolated as 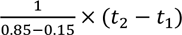 with *t*^2^ = *t*(*V* = 0.85). The *ripening relative speed* s is computed as 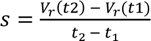. The same treatment was performed for a “mean berry”, obtained by averaging all individual raw kinetics, before computing *V*_0_, *V*_*max*_, *V*_*r*_ and *V*_*s*_. **A** Relative growth kinetics of each measured berry (grey lines), compared with the mean berry (red dotted line). Inset: histogram of initial (*V*_0_) and maximum (*V*_*max*_) volumes. **B**, **C** and **D** respectively show the timing, duration and relative speed of berry ripening, as a function of the relative volume gain (*V*_*max*_ − *V*_0_)/*V*_0_. The grey dotted horizontal line represents the mean value for the y-axis. The red stars indicate the corresponding values for the “mean berry”. In **D**, the blue dotted line corresponds to the linear regression between x and y axis.

**Fig. 9-.**
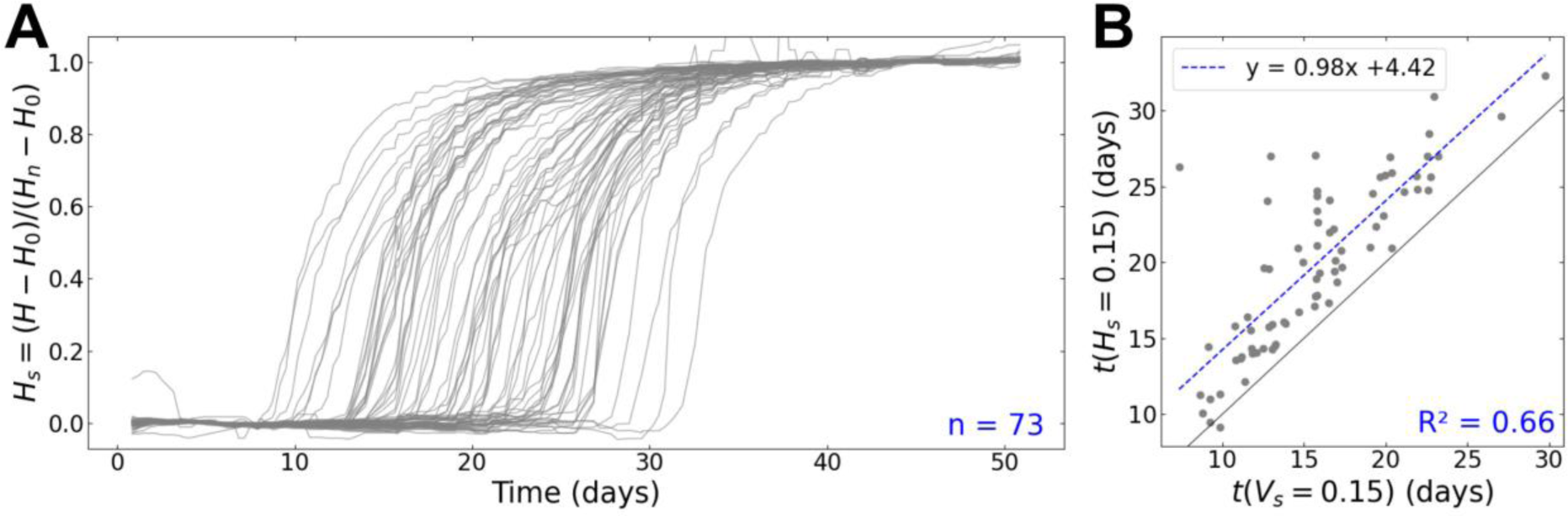
Individual coloration kinetics of berries within a grapevine bunch. Centred hue (*H*) kinetics were automatically computed for *n* = 73 berries, by applying the full image analysis pipeline on time-series of 138 images from 3 different camera views (120° difference) of the same grapevine bunch. **A** Scaled coloration kinetics of each measured berry (grey lines). **B** Relation between growth resumption time *t*(*V*_*s*_ = 0.15) and coloration time *t*(*H*_*s*_ = 0.15). The grey line is the x=y diagonal, and the blue dotted line shows the linear regression between growth resumption and coloration times.

## Discussion

### A method to quantify the variability of ripening kinetics within an asynchronous cohort of growing berries

The automated tracking of the asynchronous ripening of individual berries within one grapevine bunch allowed us to revisit the basic growth rates of ripening berries, and to propose, for the first time, an analysis of their growth and colour kinetics for a statistically significant number of observations, with an unprecedented time resolution (less than one day). In line with preliminary reports on other cultivars, we found that the doubling time of individual berry volume does not exceed three weeks, which differs from the 32 to 56 days duration of sugar loading reported in a panel of 36 international cultivars, following random sampling and averaging 50 berries over time [Suter et al., 2021]. This confirms that by neglecting berry asynchrony as a confounding variable, the ripening time was roughly two fold overestimated so far. This has also strong impacts on deconvolving the effects of annual variations in light, temperature and rainfalls on growth and sugar loading intensities. The gap with single fruit kinetics appears considerable, and it should be noted that the intra-bunch variability documented here far exceeds the phenological and compositional drifts observed over the last half-century, as consequences of climate change [Bécart et al., 2022]. It is therefore likely that minor phenological changes affecting the population age pyramid may have been previously misinterpreted as kinetic or metabolic changes intrinsic to the ripening process. It therefore becomes urgent to better document which part of the GxE interaction is mediated by the temporal structure of the population and the fundamentally different part linked to metabolic variation during berry ripening.

### A generic method to infer the shape of partially hidden fruits on an image

While numerous computer vision approaches have been developed to identify and measure fruits using deep-learning [Ganesh et al., 2019; Gonzalez et al., 2019; Ni et al., 2021; Jia et al., 2020; Gené-Mola et al., 2020; Perez-Borrero et al., 2021; Shen et al., 2022], they mostly aim at inferring their occlusion boundaries (i.e. visible edges), which differs from the true contours of the object of interest in the case of overlapping fruits, thus preventing to access their actual size. Instead, our segmentation method was designed to infer the non-visible part of overlapping fruit shapes, directly in the deep-learning process. The original and fast annotation strategy we introduced allowed to implicitly constrain the model during training to produce elliptical masks, without the need to pass by a long annotation of image edges, or to make this constrain explicit in the model architecture, as in Ellipse R-CNN [Dong et al., 2021]. The counterpart of this strategy is to restrict the inference to sufficiently visible berries, which we managed to do by training a model at detecting only berries with more than 50% visible contours. Using such a binary criterion can lead to ambiguities during both annotation and prediction, but our results suggest that it does not degrade the biological outputs. Still, including this filter in the detection step does not allow to detect all visible berries, which would be a limit for counting. An alternative would be to first use a more exhaustive fruit detector, and delegate the task of filtering measurable berries to a classifier. This would allow for classical counting strategies (e.g. [Zabawa et al., 2020]) to be combined with our physiological measurements in a single pipeline. While our method was only evaluated in controlled conditions within the PhenoArch platform, it could probably be adapted to other environments such as field conditions, at the cost of retraining the model with newly labelled data. This generalisation will be facilitated by our open-source implementation that allows re-use, and by our fast and robust annotation strategy (about 100-150 berries per hour), compared to traditional approaches that rely on annotating visible edges. Although this study focuses on grapevines, we think that our method could be applied to any other fruit that can be approximated to have an ellipsoidal shape.

### Berry tracking could be extended to other conditions, assuming a stable image acquisition setup

This work was carried out on images from a high-throughput phenotyping platform, where controlled conditions and standardised image acquisition facilitated the temporal tracking of individual berries. While we adapted the tracking algorithm to better tolerate slight movements of the bunch and camera, our results still showed that the tracking performance can be improved by almost 50% by stabilising the image acquisition. In particular, tracking performance may greatly improve by avoiding non-linear relative movements (e.g. rotations) of the bunch and the camera, preventing deformations inside the bunch, and keeping the entire bunch in the camera’s field of view. While more advanced image analysis methods may address these issues in the future, we argue that it is more effective to avoid such situations during image acquisition. Still, our method allows to visualize the aforementioned discontinuities in the image time-series via the computation of a distance matrix (Fig. 3C). This could allow to improve the tracking performance in a semi-automatic way, by re-running the automatic tracking for all periods delineated by discontinuities, and then manually mapping berry labels at each discontinuity. Lastly, choosing the right image acquisition timings is essential for subsequent analysis. Indeed, quantifying rapid dynamics such as berry colour changes needs a sufficiently high frequency of image capture, and the standardisation of ripening dynamics requires including both their initial and final plateaus during the observation period.

## Conclusion

We introduce a fully-automatic open-source method to detect, segment and track overlapping berries in a time-series of grapevine bunch images. This non-destructive method gives direct access to the growth and colour kinetics of individual berries within a bunch. Coupled with high frequency image capture, this makes it possible to quantify undocumented aspects of individual fruit development, and to characterise their asynchrony at the population level. Using this method in real time during future experiments could allow the design of new sampling strategies that will consider the bunch as a population of unsynchronized berries, rather than an ideal, average berry, and lead to a complete revisitation of the ripening dynamics. In particular, the GxE effects on harvest and fruit quality will be clearly attributed not only to physiological changes in the ripening process, but also to changes in the age structure of the whole population of berries. The complete automation of our method is also fully compatible with high-throughput phenotyping, providing the opportunity to study these detailed GxE interactions on physiology and asynchrony of berry ripening for large plant panels.

## Declarations

### Ethical Approval

Not applicable.

### Authors’ contributions

LCB supervised the experiment and acquired the data. BD and MC tested and compared different image analysis pipelines. MC provided the first proof of concept on grapevine in natural conditions. BD and CF designed the final pipeline and analysed the resulting data. LCB and CR provided advice on the conception of the pipeline. BD and CR wrote the manuscript and CF, LCB, CR, TS reviewed and edited it. All the authors have approved the manuscript and have made all required statements and declarations.

### Availability of data and materials

The source code for both training and prediction, notebook examples and trained model are available on Github (github.com/openalea/deepberry) under an Open Source licence (Cecill-C).

## Acknowledgements

We are grateful to all members at the M3P platforms for providing technical support, conducting the experiments and collecting data.

## Consent for publication

Not applicable.

## Additional Files

**Additional File 1 Video (.mp4) of berry segmentation, detection and tracking outputs for 3 grapevine bunches.** For each plant, the video displays a time-series of 62 to 66 labelled segmented RGB images, obtained after running the full berry segmentation, detection and tracking pipeline. Raw images were captured with a median interval of 8h. Each colour corresponds to one tracking label. Segmented berries without labels are drawn as white empty ellipses. t indicates the order of each image in the time-series.

**Additional File 2 - Growth and coloration kinetics of several individual grapevine berries.** Repetition of the results shown in Fig. 7 for more berries. Each subplot displays the Volume (mL) (**A**) or Centred hue (deg) (**B**) measured over time (days) on an individual berry, after running the full image analysis pipeline on a time-series of 138 images, from 3 different camera views (120° difference) of the same grapevine bunch. All points are coloured using the corresponding average hue values. In **A**, the red curve corresponds to a 8-days moving median smoothing. In **B**, the grey area corresponds to the standard deviation of the centred hue value observed within the berry segmentation mask.

**Additional File 3 Analysis of berry detection errors in the test subset.** Analysis of the False Positive (FP) and False Negative (FN) errors found when comparing berries detected by the pipeline to manually annotated berries, on the grapevine bunch images from the test subset. **A** Manual classification of detection errors as pea-sized berries, non-small (i.e. not pea-sized) berries, and non-berry objects. Non-small berries are further classified according to their percentage of visible contours (ct). **B** Distribution of detected berry sizes after segmentation, for all berries (top subplot), FP (middle subplot) and FN (bottom subplot). n: number of detected berries.

**Additional File 4 Analysis of abrupt transitions in time-series of grapevine bunch images. A** Heat map of the distance matrices obtained after tracking berries in time-series of 138 grapevine bunch images, for 6 different plants. Vertical red lines correspond to the empiric annotation of time-steps exhibiting abrupt transitions in these matrices. **B** Tracking coverage (*T*_*c*_) over time obtained for these time-series. The dashed blue vertical line represents the time *t*_*root*_ used to initialise the tracking.

