## Supplementary figures and images for "Ripening dynamics revisited: an automated method to track the development of asynchronous berries on time-lapse images"

### Additional File 2

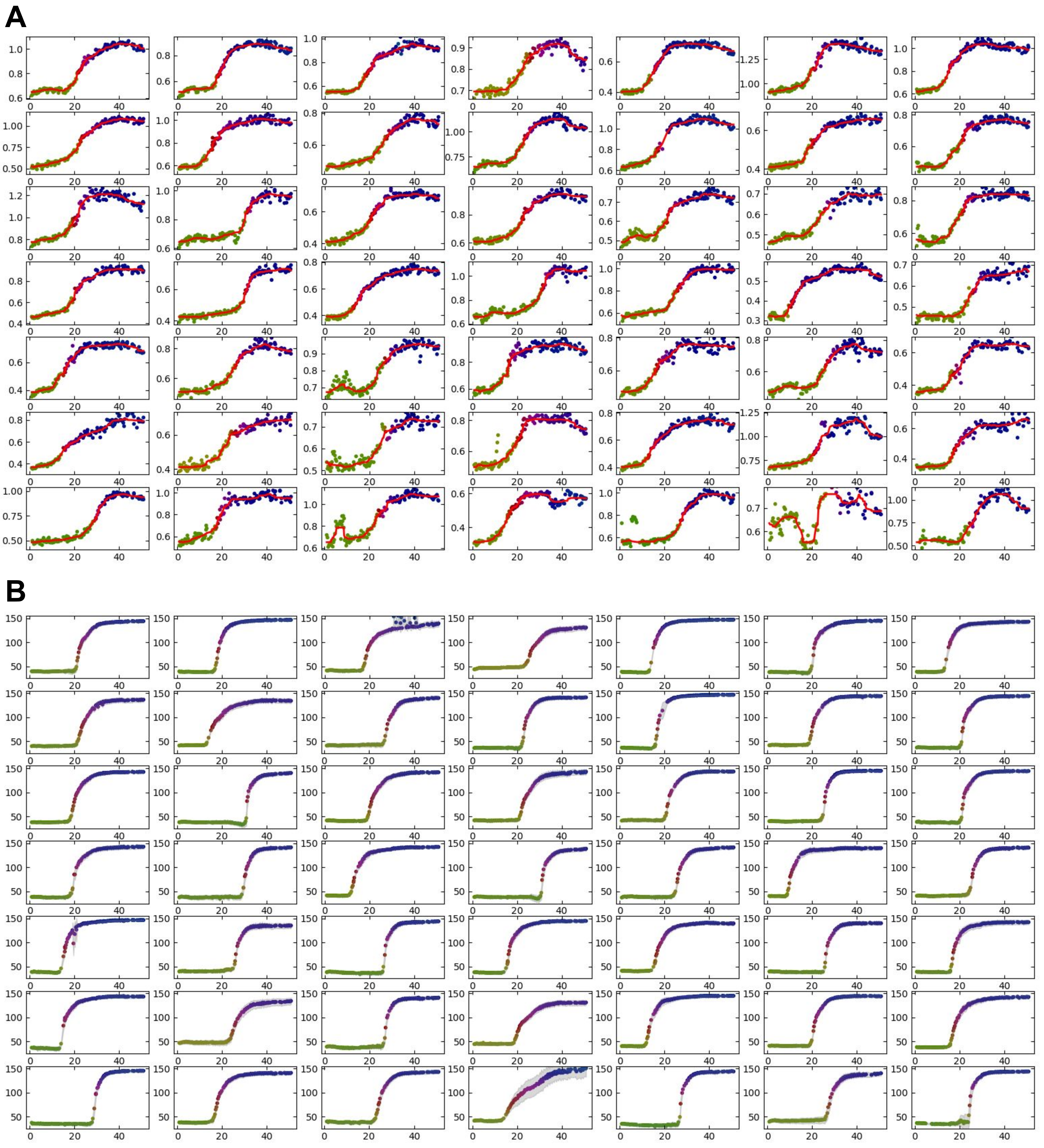

### Additional File 4

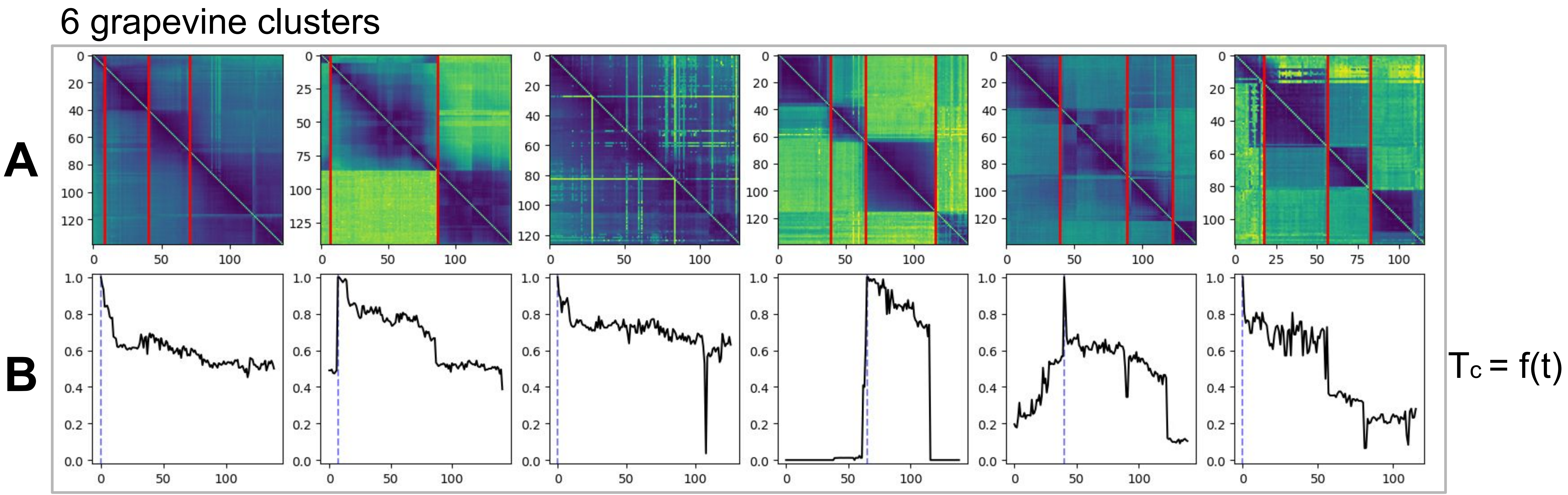
