## Additional File 3 for "Ripening dynamics revisited: an automated method to track the development of asynchronous berries on time-lapse images"

ct = % of visible berry contours

A

| | Pea-sized berry | Non-small berry ct = (50 ±10) % | Non-small berry ct < (50 ±10) % | Non-small berry ct > (50 ±10) % | Not a berry | Total |
|---|---|---|---|---|---|---|
| FP | 3% 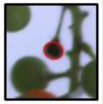 | 56% 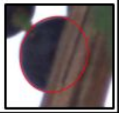 | 20% 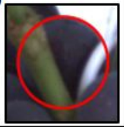 | (always labeled) | 23% 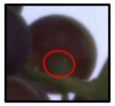 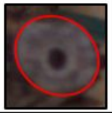 | 100% (n=64) |
| FN | 27% 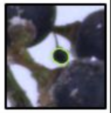 | 25% 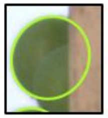 | (never labeled) | 47% 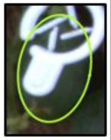 | (never labeled) | 100% (n=109) |

B

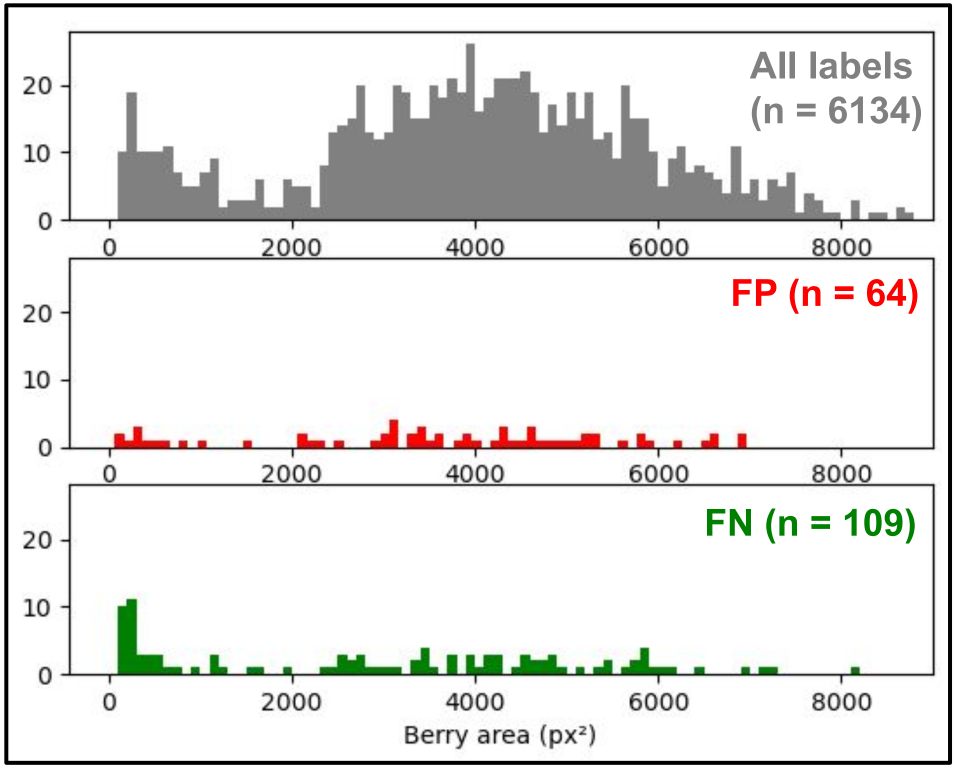
